# Theta-mediated conceptual reinstatement in vmPFC precedes perceptual reinstatement in ventral visual cortex during memory recall

**DOI:** 10.64898/2025.12.17.695049

**Authors:** Lei Zhang, Mohan Yuan, Asaf Gilboa, Claude Alain

**Author notes:** **Correspondence** Lei Zhang, Ph.D. 3560 Bathurst St, Toronto, Ontario, Canada M6A 2E1.

## Abstract

When recalling past episodes, different features of an experience, such as conceptual meaning and perceptual detail, are reconstructed and reinstated across distributed cortical regions. However, current models of human memory remain unclear how these feature-specific reinstatements unfold over time, and whether they interact hierarchically during memory retrieval. Using magnetoencephalography (MEG) and time-resolved encoding-retrieval cross-phase classification analysis, we compared the time course of conceptual and perceptual reinstatement during cued visual recall. Conceptual information decoding in the ventromedial prefrontal cortex (vmPFC) preceded perceptual information decoding in the ventral visual cortex (VVC), and the two were expressed in distinct frequency bands: theta (4–8 Hz) for conceptual and gamma (30–40 Hz) for perceptual information. Cross-correlation and spectral Granger causality analyses revealed that during reinstatement conceptual information in vmPFC predicted and directed perceptual information in VVC, with theta-band information flow predominantly from vmPFC to VVC. In addition, bottom-up connectivity from VVC to vmPFC was expressed in the alpha band. These findings suggest that memory retrieval proceeds from abstract conceptual reconstruction to the reinstatement of perceptual details, mediated by theta oscillatory communication between prefrontal and sensory areas. We propose a hierarchical interactive model in which vmPFC initiates conceptual activation through theta-band top-down signals, while VVC provides alpha-band feedback for evaluating reconstructed perceptual details.

## Introduction

When you recall a beautiful object that fascinated you yesterday, you may first remember that it was a flower and then gradually fill in perceptual details such as its color and shape^1–3^. This hierarchical reinstatement process during memory recall aligns with theories that describe memory engrams as distributed across multiple, functionally connected cortical regions, with each region corresponding to distinct features or aspects of the memoranda being stored^4–6^. This organization enables different components of memory, such as perceptual, contextual, and conceptual details, to be reactivated within distributed neural ensembles that are functionally interconnected as part of a unified engram complex^7^. Current models of memory suggest detailed perceptual features are represented in posterior associative modal neocortical regions, whereas abstract conceptual features are represented in more anterior a-modal regions such as the ventromedial prefrontal cortex (vmPFC)^8–11^ (but see^12^). However, current models of human memory are unclear regarding the time-course of memory reinstatement and directionality of interactions between perceptual representations in the posterior neocortex and conceptual representations in the vmPFC during episodic memory retrieval.

Cortical reinstatement, which refers to the reactivation of cortical activity patterns present during initial encoding, is considered a core neural mechanism of episodic memory retrieval^12–19^. Functional MRI studies have shown that recalling visual scenes reactivates the same regions engaged during picture encoding and reinstates the fine-grained spatial patterns observed at encoding, particularly within ventral visual areas (VVC) when the recalled stimuli are complex, real-world pictures^14,16,20,21^.

Moreover, the strength of reinstatement in sensory cortices predicts the vividness of recollected visual memories^13,16,20–23^. Converging neuroimaging and neuropsychological evidence also suggests that higher-order prefrontal regions, particularly the vmPFC, are involved in the retrieval of conceptual representations^9,24–26^. Together, these findings suggest that perceptual and conceptual reinstatement reflect distinct yet complementary aspects of memory retrieval. The reconstructive (rather than reduplicative) nature of episodic memory may depend on reducing dimensionality at encoding and expanding memory codes at retrieval (dimensionality transformations)^27,28^. This suggests that the reconstruction of vivid perceptual details in the posterior cortex entails an expansion of stored compressed conceptual representations in medial prefrontal cortex^28,29^;therefore, conceptual representations should precede perceptual reinstatement^30,31^.

While functional magnetic resonance imaging (MRI) studies have characterized the location of reinstatement, electrophysiological techniques such as magnetoencephalography (MEG) enables tracking of both where and when reinstatement unfolds with millisecond precision. Encoding and retrieval cross-phase multivoxel pattern analysis (MVPA) can be used to quantify how closely voxel-wise activation patterns during retrieval resemble those recorded during encoding, providing a direct measure of reinstatement of the original memory trace^14,20^. Combining MEG recordings with time-resolved source MVPA provides a unique opportunity to examine the temporal dynamics of cortical reinstatement. In the present study, we used this approach to directly compare the time courses of perceptual reinstatement in the VVC and conceptual reinstatement in the vmPFC during memory retrieval.

Recent behavioral and EEG evidence supports the idea that conceptual information is accessed earlier than perceptual details during memory retrieval^30,31^, consistent with the idea of dimensionality expansion^27^. Specifically, high-level conceptual features can be accessed more rapidly than low-level perceptual features during recall of visual objects, which is the reverse of the sequence typically observed during encoding^30,31^. Consistent with this behavioral pattern, an EEG study has shown that neural activity associated with conceptual features reemerges earlier than activity linked to perceptual features^31^. Building on these findings, we hypothesize that during visual memory retrieval, conceptual reinstatement in the vmPFC would occur earlier than perceptual reinstatement in the VVC.

Evidence from electrophysiological studies suggests that theta oscillations (4–8 Hz) play a key role in episodic memory retrieval^32–36^. For instance, theta activity has been linked to successful memory recall and reinstatement of past experiences^33–35^. Theta rhythms may enable the long-range transfer and integration of mnemonic information between the hippocampus and neocortical regions^33,34,37,38^, an interaction that is causally linked to vivid re-experiencing of episodic memory^39^. Previous studies have shown that theta coherence increases between the vmPFC and sensory association areas during retrieval, suggesting that theta oscillations mediate the exchange of information along this prefrontal–sensory pathway^33,40–42^. Evidence from human and animal studies is consistent with the vmPFC exerting top-down control over reinstatement in the sensory areas, biasing or initiating retrieval to align with prior knowledge and goals^43–46^. This long-range synchronization is believed to convey mnemonic information across relevant networks, enabling the vmPFC to integrate conceptual representations and guide reinstatement of perceptual details in the VVC. Based on this framework, we hypothesized that theta oscillations would index the interaction between vmPFC and VVC during visual memory recall.

To test these hypotheses, we recorded neuromagnetic activity during a cross-modal automatic cued-retrieval paradigm that maximized the time-lock accuracy of item recall and minimized confounding perceptual processing and memory strength. Combining this paradigm with time-resolved multivoxel pattern analysis allowed us to examine the timing, frequency, and direction of conceptual and perceptual reinstatement with high temporal precision. We predicted that 1) conceptual reinstatement in the vmPFC would occur earlier than perceptual reinstatement in the VVC, 2) that conceptual reinstatement in vmPFC would correlate with and predict later perceptual reinstatement in VVC, and 3) that theta oscillations would convey information from vmPFC to VVC during recall.

## Results

### Conceptual reinstatement showed an advantage in vmPFC, while perceptual reinstatement showed an advantage in VVC

Participants learned four audiovisual pairings that crossed conceptual (architecture vs. plants images; human vs. tool sounds) and perceptual (color vs. black-and-white; low-vs. high-frequency) features (Figure 1). Before MEG recording, they completed extensive training to ensure the sounds would obligatorily and immediately lead to recall of the associated item. During MEG recording, participants completed blocks that began with re-encoding of all four audiovisual stimuli, followed by cued-recall trials in which either a sound cued the recall of its paired picture or a picture cued the recall of its paired sound. Each recalled item was followed by a judgment of vividness. Here, we test the hypothesis of format transformation during visual object episodic reactivation; therefore, we only analyze auditory-cued picture recall trials.

**Figure 1.**
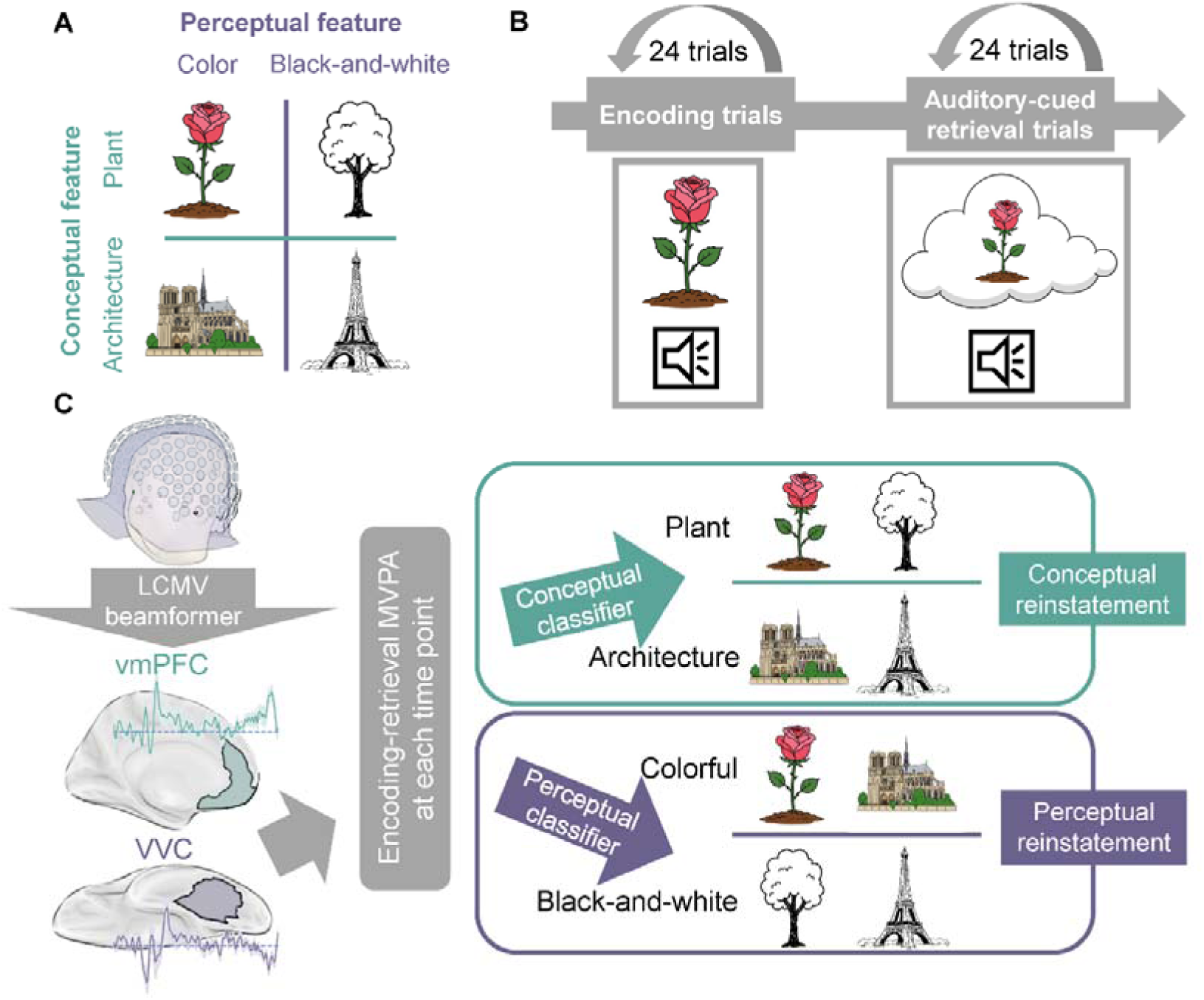
Experimental paradigm and analysis pipeline. (A) Four picture stimuli used in the experiment were organized along two orthogonal dimensions: a perceptual dimension (color vs. black-and-white) and a conceptual dimension (architecture vs. plant). (B) MEG run structure. Each run began with twenty-four encoding trials in which participants viewed intact audiovisual pairs. This was followed by twenty-four cued-recall trials, including auditory-cued picture retrieval and picture-cued sound retrieval. (C) After MEG data acquisition, sensor-level data were source-localized using an LCMV beamformer. We then extracted trial-wise time series from vmPFC and VVC regions of interest. Encoding–retrieval MVPA was performed at each time point to quantify perceptual and conceptual reinstatement. vmPFC, ventromedial prefrontal cortex; VVC, ventral visual cortex; MVPA, multivariate pattern analysis.

First, we compared the reinstatement of conceptual and perceptual features time series separately in vmPFC and VVC. Reinstatement strength was quantified with encoding–retrieval cross-phase classification accuracy at each time point. We trained the conceptual and perceptual classifiers using encoding phase data and tested them using recall phase data (for details, see the Methods section). Higher classification accuracy denotes more robust evidence for reinstatement of a feature type.

We found that although both features could be decoded in both regions, reinstatement of conceptual features was stronger than reinstatement of perceptual features in vmPFC from 110 ms to 160 ms (Fig. 2A, p_fwe_ < 0.05), while perceptual feature reinstatement was stronger than conceptual feature reinstatement in VVC from 190 ms to 230 ms (Fig. 2B, p_fwe_ < 0.05). This supports models predicting that conceptual features are more prominently represented in the vmPFC during object recall and perceptual features are more prominently represented in posterior visual association areas^8,47^.

**Figure 2.**
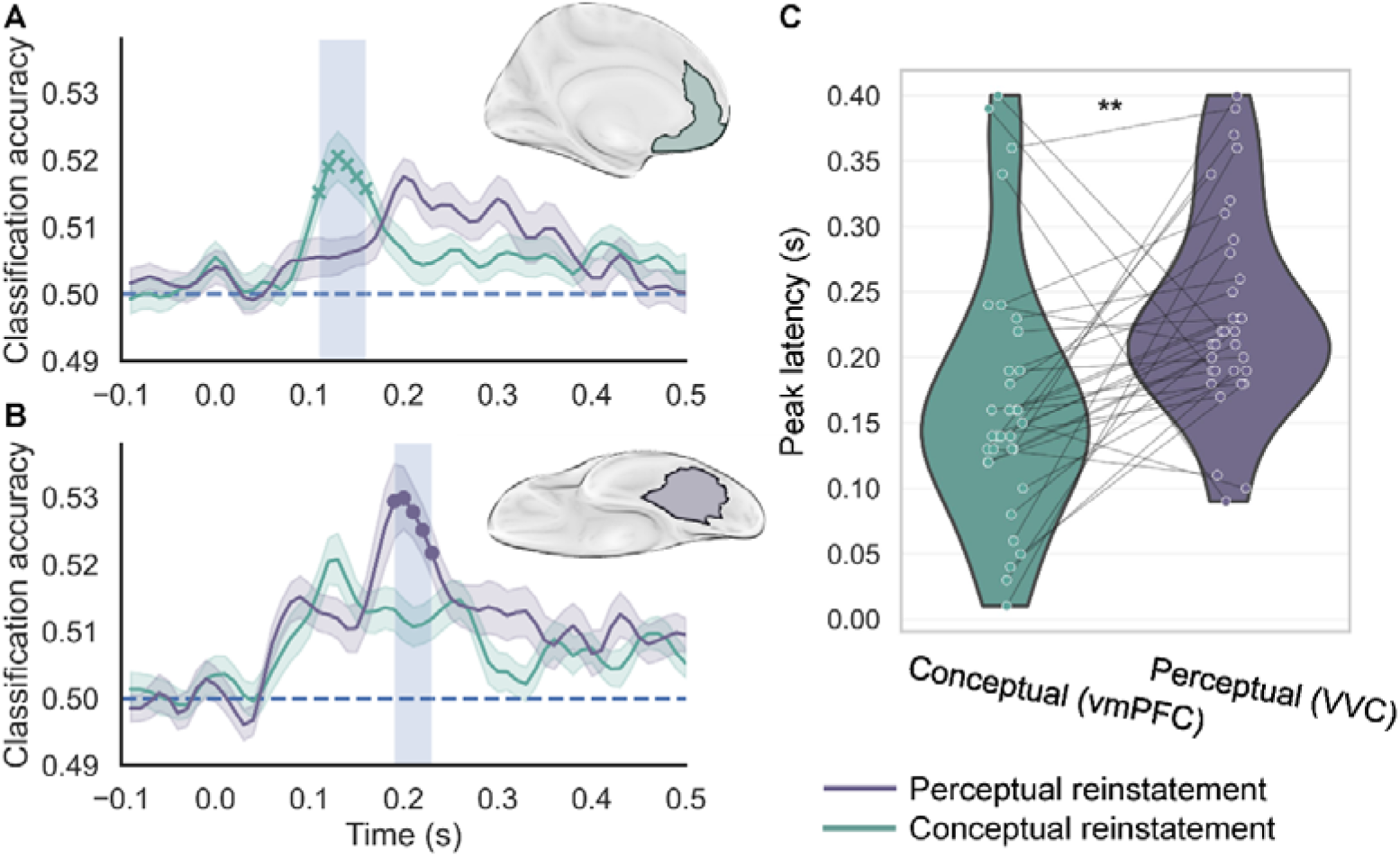
Conceptual reinstatement in vmPFC precedes perceptual reinstatement in VVC. (A). vmPFC showed stronger conceptual reinstatement than perceptual reinstatement from 110-160 ms (p_fwe_ < 0.05). (B) VVC showed stronger perceptual reinstatement than conceptual feature reinstatement from 190-230 ms (p_fwe_ < 0.05). (C) The onset latency of conceptual reinstatement in vmPFC was significantly earlier than the onset of perceptual reinstatement in VVC (p_fwe_ < 0.05). vmPFC, ventromedial prefrontal cortex; VVC, ventral visual cortex; **, p < 0.01.

### Conceptual reinstatement in vmPFC precedes perceptual reinstatement in VVC

As shown in Figures 2A and 2B, we observe that the conceptual advantage in vmPFC and the perceptual advantage in VVC appear in distinct time windows. We extracted the peak latency of conceptual reinstatement in vmPFC and the peak latency of perceptual reinstatement in VVC for each participant to directly test whether conceptual reinstatement in vmPFC preceded perceptual reinstatement in VVC. We found that conceptual reinstatement in vmPFC was significantly earlier than perceptual reinstatement in VVC (Fig. 2C, average latency of conceptual reinstatement in vmPFC: 165ms, average latency of perceptual reinstatement in VVC: 234ms, paired two-tailed t-test: t(32) = −3.052, p = 0.005). These results support our first hypothesis that conceptual reinstatement in vmPFC precedes perceptual reinstatement in VVC.

### Conceptual advantage in vmPFC is expressed in the theta band, while leads perceptual advantage in VVC is expressed in the gamma band

Next, we asked whether conceptual advantage in vmPFC and perceptual advantage in VVC are expressed in different frequency bands. To examine the frequency-specific effect of the conceptual advantage and perceptual advantage, we first filtered the broadband neural signal into multiple narrowband signals and then performed the same encoding-retrieval cross-phase classification analysis on each narrowband signal (for details, see the Methods section). For the statistical analyses, we averaged across four leading frequency bands (theta: 4–8 Hz, alpha: 8–13 Hz, beta: 13–30 Hz, gamma: 30–40 Hz), and compared the time series of conceptual and perceptual reinstatement separately in the vmPFC and VVC across these frequency bands.

In the vmPFC, we found that conceptual reinstatement only showed an advantage over perceptual reinstatement in the theta band from 0 ms to 70 ms and from 110 ms to 160 ms (Fig. 3A, p_fwe_ < 0.05), with no effects in other bands (Fig. S1A, all p_fwe_ > 0.05). Meanwhile, in VVC, perceptual reinstatement showed an advantage over conceptual reinstatement in the gamma band from 300 ms to 370 ms (Fig. 3B, p_fwe_ < 0.05), with no difference in other bands (Fig. S1B, all p_fwe_ > 0.05). These results indicate that conceptual and perceptual reinstatement are associated with distinct frequency bands, with the conceptual advantage in the vmPFC expressed in the theta band and the perceptual advantage in VVC reflected in the gamma band.

**Figure 3.**
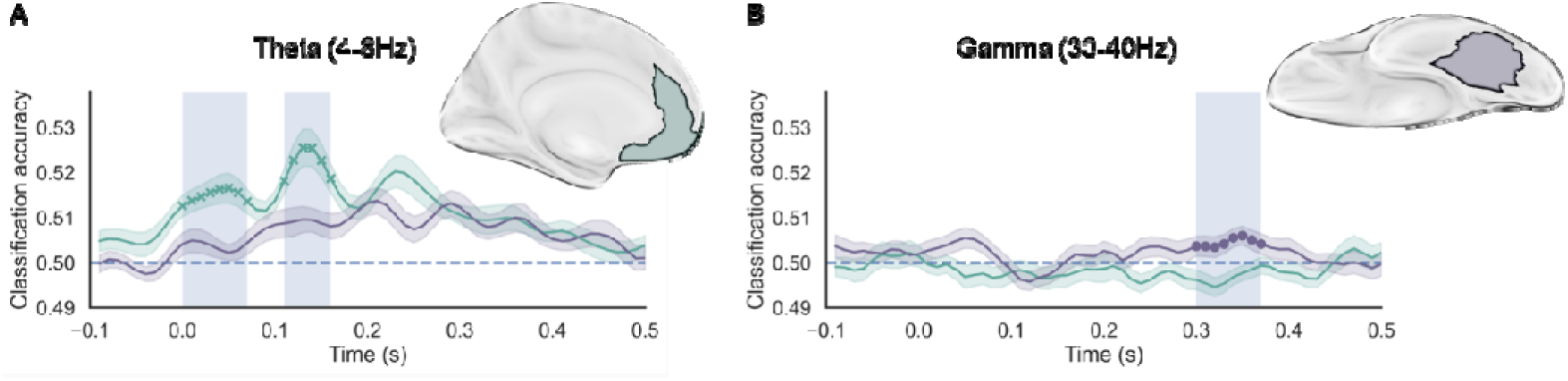
Conceptual advantage in vmPFC expressed in theta band while leads perceptual advantage in VVC expressed in gamma band. (A). In vmPFC, conceptual reinstatement was significantly stronger than perceptual reinstatement in the theta band during the two time windows: 0–70 ms and 110–160 ms (< 0.05). (B) In contrast, VVC showed significantly stronger perceptual than conceptual reinstatement in the gamma band from 300–370 ms (< 0.05). vmPFC, ventromedial prefrontal cortex; VVC, ventral visual cortex.

### Conceptual reinstatement in vmPFC directly influenced perceptual reinstatement in VVC

Based on the finding that conceptual reinstatement in vmPFC preceded perceptual reinstatement in VVC, we hypothesized that conceptual reinstatement in vmPFC directly influenced perceptual reinstatement in VVC. We performed informational cross-correlation analysis, measuring the similarity between the two time series or signals as a function of the time lag between them from 10 ms to 150 ms in 10 ms steps. This analysis allows for the examination of the directed correlation between two reinstatement time series.

We compared the correlation coefficients at each time lag k between r_conceptual→perceptual_(k) (conceptual reinstatement precedes perceptual reinstatement) and r_perceptual→conceptual_(k) (perceptual reinstatement precedes conceptual reinstatement) with cluster-based correction. This comparison indicated that ^r^_conceptual→perceptual_ ^was significantly greater than r^_perceptual→conceptual_ ^in time lags^ ranging from 70 to 90 ms (Fig. 4A, p_fwe_ > 0.05). Next, we extracted the peak conceptual classification accuracy in vmPFC and the peak perceptual classification accuracy in VVC, each identified at subject-specific peak latencies. We found that stronger conceptual reinstatement was significantly linked to later perceptual reinstatement (Fig. 4B, Pearson correlation r = 0.36; p = 0.039). These results support our hypothesis that conceptual reinstatement in vmPFC predicts perceptual reinstatement in VVC.

**Figure 4.**
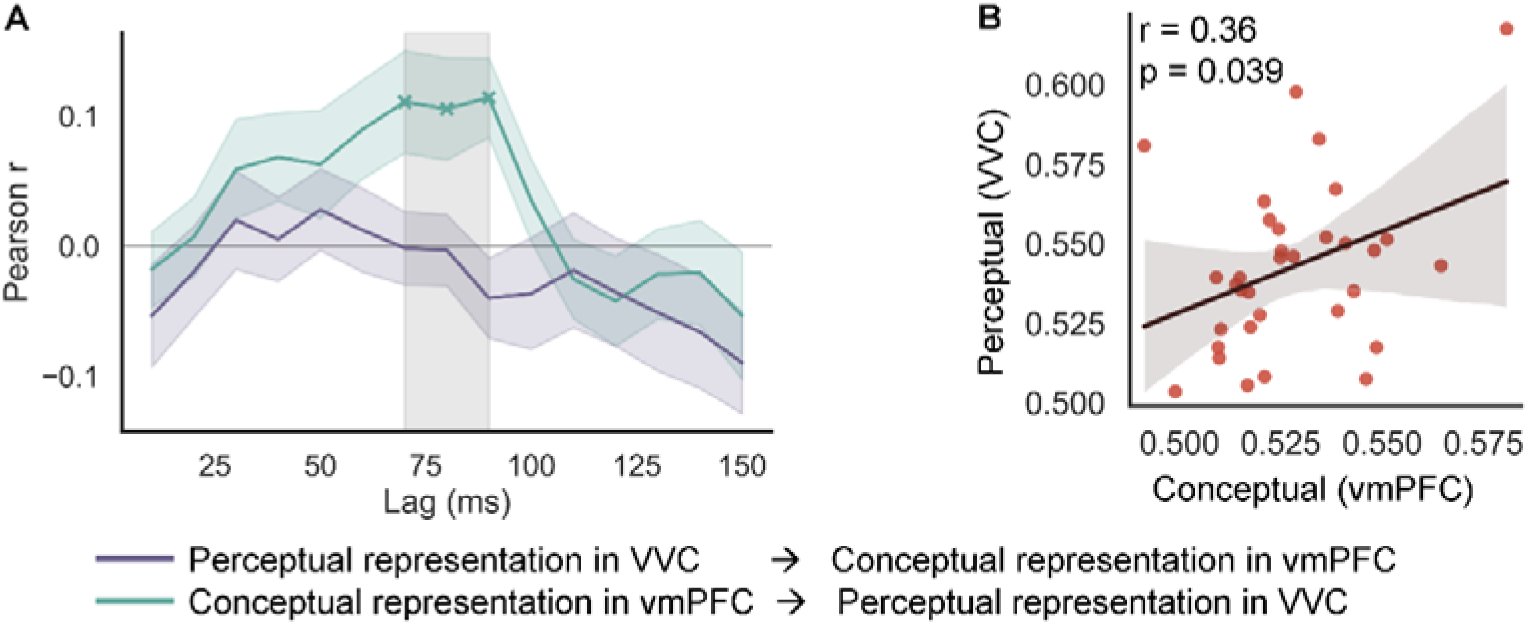
Conceptual reinstatement in vmPFC directly influenced perceptual reinstatement in VVC. (A) vmPFC conceptual reinstatement exerted a direct influence on VVC perceptual reinstatement during the 70–90 ms time window. (B). Stronger conceptual reinstatement could significantly predict later perceptual reinstatement. vmPFC, ventromedial prefrontal cortex; VVC, ventral visual cortex.

### vmPFC drives VVC activity during retrieval via theta-band oscillations

Lastly, we tested the third hypothesis that theta frequency plays an important role in conveying information from vmPFC to VVC using spectral state-space Granger causality (GC) analysis. We computed the spectral GC estimate from vmPFC to VVC and from VVC to vmPFC across frequencies ranging from 2 to 40 Hz using trial-wise neural activities during cued memory recall. As in the frequency-specific encoding-retrieval cross-phase classification analysis, we also averaged the spectral GC estimates across four main frequency bands (theta: 4–8 Hz, alpha: 8–13 Hz, beta: 13–30 Hz, and gamma: 30–40 Hz). Then, we compared the spectral GC estimate from vmPFC to VVC with the spectral GC estimate from VVC to vmPFC within each frequency band.

We found that the spectral GC estimate from vmPFC to VVC was significantly stronger than the estimate from VVC to vmPFC in theta neural oscillation (Fig. 5, t(32) = 2.89, p = 0.007). This indicates that information flow is predominantly from vmPFC to VVC in the theta band during cued picture retrieval. These findings support our fourth hypothesis that information flow from vmPFC to VVC is mediated by theta oscillations during picture memory recall. Unexpectedly, we found that the spectral GC estimate from VVC to vmPFC exceeded the spectral GC estimate from vmPFC to VVC in alpha oscillation (Fig. 5, t(32) = −2.71, p = 0.011). In summary, during visual memory retrieval, vmPFC and VVC interact bidirectionally, with coupling from vmPFC to VVC emphasized in the theta band and coupling from VVC to vmPFC emphasized in the alpha band.

**Figure 5.**
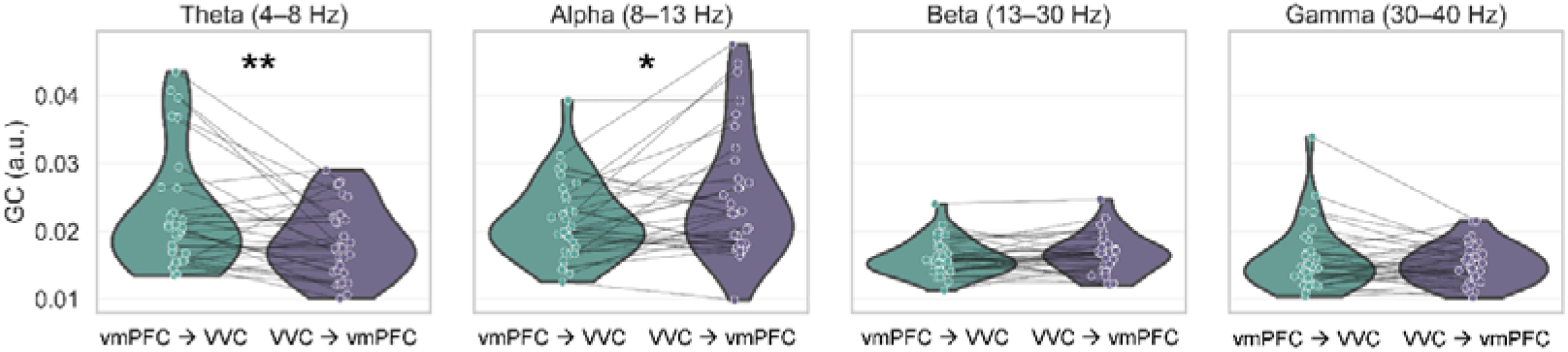
Bi-directional connectivity between vmPFC and VVC. vmPFC drives VVC activity via theta-band top-down influences during retrieval, whereas VVC provides bottom-up feedback to vmPFC through alpha-band connectivity. vmPFC, ventromedial prefrontal cortex; VVC, ventral visual cortex; *, p < 0.05; **, p < 0.01.

## Discussion

Our study provides converging evidence that conceptual and perceptual reinstatement during visual memory recall are temporally, spectrally and spatially dissociable yet functionally and directionally interdependent. Using time-resolved MEG source encoding-retrieval cross-phase classification analysis, we found that conceptual reinstatement in the vmPFC preceded perceptual reinstatement in the VVC, that these reinstatements were expressed in distinct frequency bands (theta oscillation: conceptual reinstatement in the vmPFC, and gamma oscillation: perceptual reinstatement in the VVC, respectively), and that information flowed predominantly from vmPFC to VVC through theta-band oscillations. Together, these findings support the view that memory retrieval proceeds from abstract conceptual reconstruction to the reactivation of perceptual detail, mediated by theta oscillatory communication between vmPFC and sensory areas.

The temporal precedence of vmPFC over VVC supports the notion that memory retrieval is initiated by the reinstatement of high-level conceptual information that subsequently guides perceptual reconstruction. This finding directly supports and extends the current models, including Trace Transformation Theory (TTT), which proposes that conceptual and perceptual information are stored as complementary traces in anterior and posterior cortical systems, respectively^8,9,47–49^. Extending these models, our data demonstrate how different formats of these memory traces reflecting different features are dynamically coordinated in both the time and frequency domains. Furthermore, conceptual reinstatement in vmPFC may thus provide a top-down scaffold that constrains and refines perceptual reconstruction in VVC. This hierarchical temporal structure refines the models of episodic retrieval by showing that conceptual information is encoded as a distinct memory trace early^7^ in retrieval, rather than extracted over time^47,49^, and that its reinstatement actively drives the reactivation of perceptual representations rather than passively co-occurring with them.

The temporal difference observed in this study is consistent with previous behavioral and EEG findings showing that conceptual features are retrieved faster than perceptual features^30,31^. This suggests that even within the same object, features at different representational levels vary in their accessibility during retrieval, consistent with models that suggest dimensionality transformation during the encoding and retrieval of episodic memory^27,30,31^. These different codes are represented in distinct cortical regions, with conceptual information in the anterior vmPFC and perceptual information in the posterior sensory cortices^8,9^. Here, we show a mechanism by which representations at different hierarchical levels could interact during memory retrieval.

Perceptual-to-conceptual transformation is an important process that helps encode and store information into long-term memory^27,30,50–52^. It can reduce computational load by compressing complex sensory input to simplified storable codes, thus maximizing storage capacity. Dimensionality reduction also allows us to generalize across experiences and integrate new information into existing knowledge structures^27,50^. Our findings, consistent with previous research^30,31^, suggest that memory retrieval operates in the reverse direction of this process, namely as conceptual-to-perceptual reinstatement. The informational cross-correlation results indicated that during recall, abstract conceptual traces are first reactivated within the vmPFC, guiding the reinstatement of perceptual details in the VVC. This interpretation is further supported by the correlation analysis, which shows that stronger conceptual reinstatement in an earlier time window predicted more robust later perceptual reinstatement. Moreover, previous electrophysiological studies with patients who have focal damage to the vmPFC suggest that oscillatory coupling between vmPFC and posterior cortices is disrupted, leading to impaired encoding of episodic memory^24^ and retrieval of autobiographical information^54^. Together, these findings provide neural evidence consistent with the TTT, indicating that episodic memory may emerge from a dynamic interplay between conceptual abstraction and perceptual reconstruction across cortical hierarchies. Future studies using methods such as transcranial magnetic stimulation are necessary to determine the causal influence of vmPFC conceptual activity on VVC perceptual reinstatement in episodic memory.

We further found that conceptual reinstatement in vmPFC was expressed in the theta band, whereas perceptual reinstatement in VVC was expressed in the gamma band. This frequency dissociation aligns with the proposal that theta and gamma rhythms serve complementary mnemonic roles^36,54,55^. Previous studies have consistently reported that increased theta power or coherence is associated with successful memory performance^33,35,55–57^. Because many of these studies employed recognition or associative memory paradigms that emphasize retrieval of gist- or schema-like information, theta oscillations may primarily support the conceptual or relational aspects of memory. In comparison, gamma oscillations are more closely related to fine-grained sensory processing and perceptual encoding^58–61^. Therefore, reconstructing perceptual details during recall may elicit gamma-band activity. Theta oscillations may index synchronization within a distributed cortical–hippocampal network to reinstate abstract relational structures, whereas gamma oscillations may locally encode and reconstruct fine-grained sensory details. Such a theta-gamma division of labor suggests that conceptual and perceptual aspects of episodic memory are maintained in distinct yet coordinated neural codes.

Cross-correlation and Granger causality analyses further revealed that conceptual reinstatement in vmPFC predicted perceptual reinstatement in VVC, with theta-band information flow predominantly from vmPFC to VVC. These results indicate a top–down control mechanism through which vmPFC may orchestrate sensory reactivation by propagating mnemonic predictions or templates that guide the reinstatement of perceptual detail. This interpretation is consistent with intracranial and MEG evidence showing enhanced theta coherence between hippocampus, vmPFC, and sensory cortices during successful recall^40–42^. Theta oscillations may thus provide the temporal framework that coordinates large-scale reinstatement across conceptual and perceptual systems.

Interestingly, we also observed feedback from VVC to the vmPFC in the alpha band, suggesting that once perceptual details are reinstated, sensory regions may provide feedback to higher-order areas for evaluation and integration. Such alpha-band feedback aligns with models of predictive coding, where alpha synchronization supports feedback signalling of reconstructed sensory evidence^62,63^.

An important limitation of both the present and previous studies is that only a single conceptual dimension (plant versus architecture in the present study, animate versus inanimate in earlier work) and a single perceptual dimension (color versus black-and-white here, photo versus drawing previously) were examined. One possible contributor to the observed retrieval-time difference is the unequal discriminability or representational distance of features within conceptual and perceptual dimensions. For instance, animate and inanimate categories may be more separable in conceptual space than photos and drawings are in perceptual space. However, it is inherently difficult to balance feature distances across conceptual and perceptual domains. Future studies should include multiple conceptual and perceptual dimensions to achieve better generalization and to determine whether retrieval timing differences persist across a broader range of feature contrasts. For example, lower-level perceptual contrasts could include different brightness or contrast levels, whereas higher-level conceptual contrasts could include contextual dimensions (e.g., indoor vs. outdoor items) or functional categories (e.g., tools vs. non-tools).

Taken together, we propose a hierarchical interactive model of episodic memory retrieval (Figure 6) in which the vmPFC initiates conceptual reinstatement through theta-mediated top-down signals that guide and constrain the subsequent gamma-mediated reinstatement of perceptual details in the VVC. In turn, the VVC feeds back information to the vmPFC through alpha oscillations, supporting the evaluation of the reconstructed object and the integration of features across representational levels. This temporal and spectral cascade bridges behavioral models of the reverse retrieval hierarchy with neurophysiological mechanisms of large-scale cortical communication^30,31^. Within this framework, the vmPFC may serve as a central hub that integrates hippocampal outputs and transmits them to sensory cortices, allowing for the reconstruction of vivid episodic experiences. Future research should investigate how the vmPFC and posterior hippocampus interact to influence the reinstatement of perceptual details in posterior sensory regions.

**Figure 6.**
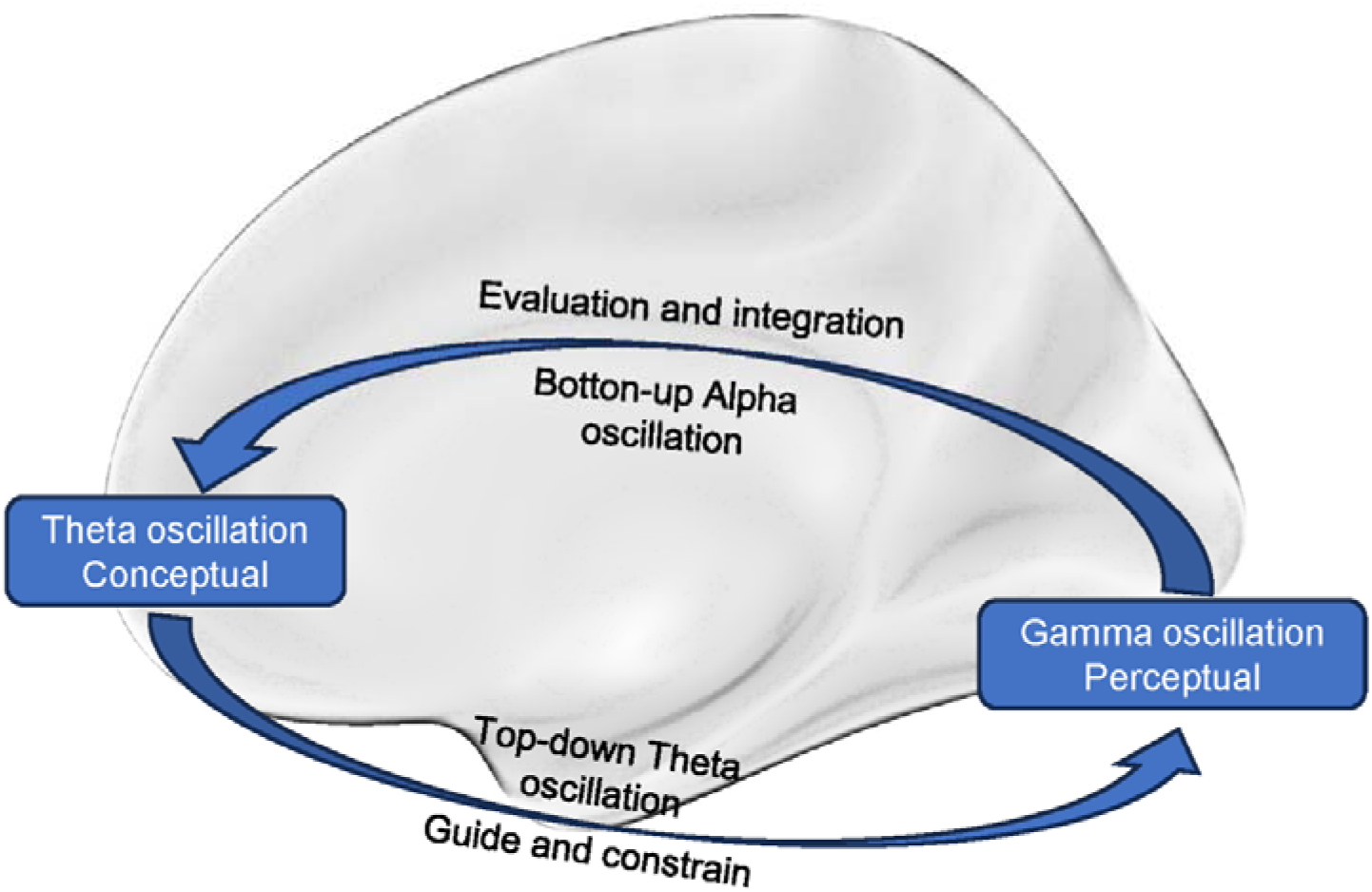
Hierarchical interactive model of episodic memory retrieval. During episodic retrieval, conceptual information is first reactivated in vmPFC, which provides high-level, schema-like constraints. This conceptual signal guides and constrains perceptual reinstatement in VVC via top-down theta-band oscillatory interactions. Meanwhile, VVC sends bottom-up alpha-band feedback to vmPFC, supplying perceptual evidence for further evaluation and integration. Together, these bidirectional interactions support the hierarchical reconstruction of vivid episodic memories. vmPFC, ventromedial prefrontal cortex; VVC, ventral visual cortex.

## Methods

### Participants

Thirty-three young adults (22.58 ± 4.54 years old, 18 female) participated in this study. Sample size was determined through a priori power analysis using G*Power version 3.1.9.7^64^. Thirty-three participants were sufficient to detect a median effect size (0.5) with 80% power using a paired, two-tailed t-test^64^. All participants were healthy right-handed individuals with no history of neurological disorder and normal hearing and vision. All participants provided written informed consent approved by the Baycrest Research Ethics Board (Approval No. 24-33).

### Stimuli

The stimuli comprised four pictures and four sounds. The four pictures and sounds had two orthogonal features, conceptual (pictures: architecture and plant pictures; sounds: human and tool sounds) and perceptual (pictures: color and black-and-white pictures; sounds: low- and high-frequency sounds). Four pictures and sounds were combined to form four paired audiovisual stimuli. The duration of the audiovisual stimuli was two seconds. Pairings were pseudorandomized for each participant, while the picture-to-sound mapping was fixed.

### Procedure

We created a cross-modal automatic-retrieval cued recall paradigm to maximize time-lock accuracy of item recall and to minimize the perception-recall process confound. Prior to the MEG scanning sessions, participants underwent a training phase to familiarize themselves with the four audiovisual stimuli. Each participant completed five training blocks. In each training block, there were twenty-four encoding trials with each audiovisual stimulus presented six times at random. Participants completed eight test trials, including four auditory-cued visual item recall and four visual-cued auditory item recall. In auditory-cued visual item recall, participants were instructed to recall the corresponding picture target as soon as the sound cue was presented, and vice versa for visual-cued auditory item recall. After five blocks, participants could recall the target automatically at cue onset, yielding high time-locking accuracy. After the training phase, participants were instructed to close their eyes and rest for about thirty minutes.

In the MEG recording session, participants completed a six-block task. In each block, twenty-four encoding trials with each audiovisual stimulus presented six times were randomly presented at the beginning. Participants were instructed to passively perceive the audiovisual pairs. Audiovisual stimuli last 2 s. Inter-trial intervals (ITI) of encoding trials were 1.0–1.5 s in 0.1 s steps. Then, there were twenty-four auditory-cued visual item recall trials and twenty-four visual-cued auditory item recall trials. Each picture target and sound target was recalled six times in each block. After recalling pictures or sounds, participants provided a vividness rating by pressing the left or right button box (left: vivid; right: not vivid). The interval between cued recall and vividness rating was 1 s. ITI of recall trials was 1.3–1.7 s in 0.2 s steps. Because comparing auditory and visual memory retrieval was not the aim of the present study, only auditory-cued picture-recall trials were analyzed.

After the MEG session, we collected participants’ high-resolution structural MRI data for source localization.

### MEG data acquisition and preprocessing

MEG data were recorded using a 275-channel CTF system (VSM MedTech) at a sampling rate of 1200 Hz. Preprocessing was performed in Python using the MNE-Python toolbox^65^. Continuous data were band-pass filtered between 1 and 120 Hz using an FIR filter. To remove artifacts, we first applied independent component analysis to the broadband sensor data. Components reflecting eye blinks, saccades, or cardiac artifacts were identified through visual inspection of component time courses and topographies and then removed. The cleaned data were subsequently epoched around stimulus onsets (−0.5 s to 2.5 s) and baseline-corrected using the −0.2 to −0.02 s pre-stimulus interval. Epochs were down-sampled to 250 Hz and manually inspected to reject residual artifacts.

### Structural data acquisition and source localization

Structural MRI data were collected on a Siemens 3T Prisma scanner with a 64-channel head coil. T1-weighted images were acquired using the magnetization-prepared rapid acquisition gradient echo (MPRAGE) sequence (TR = 2000 ms, TE = 2.85 ms, field of view = 256 × 240 mm, voxel size = 0.8 × 0.8 × 0.8 mm).

Structural T1-weighted MRIs were processed with FreeSurfer to reconstruct cortical surfaces and define the source space^66^. MEG–MRI coregistration was performed individually, run by run, by aligning fiducial points to anatomical landmarks (nasion and bilateral preauricular points). A single-shell boundary element model was then generated to model the inner-skull conductivity, from which individual forward solutions were computed for each experimental run in surface space. Linearly constrained minimum variance spatial filters were constructed for each run using empirical data (0.01–2.0 s) and baseline (−0.5 to −0.02 s) covariance matrices. Filters were built with unit-noise-gain normalization and then applied to cleaned MEG epochs to obtain single-trial source time courses. Epochs with head displacements exceeding 10 mm were excluded. Head displacement was computed from the circumcenter of three localization coils per epoch. Source estimates were cropped to −0.4–2.2 s, band-pass filtered (1–40 Hz), and down-sampled to 100 Hz. All source data were computed in native space and parcellated according to the Destrieux cortical atlas (aparc.a2009s) for subsequent ROI-based decoding analyses^67^.

### Encoding-retrieval cross-phase classification analysis

We quantified reinstatement between perception and retrieval using a ROI-wise, time-resolved cross-phase decoding analysis on source estimates. Based on our hypotheses, vmPFC ROIs included left and right G_subcallosal, G_orbital, G_rectus, S_suborbital, G_and_S_cingul-Ant, G_and_S_transv_frontopol, G_and_S_frontomargin and VVC ROIs included left and right G_oc-temp_lat-fusiform, S_oc-temp_lat,S_oc-temp_med_and_Lingual, as defined in the Destrieux (aparc.a2009s) atlas. For each participant, single-trial LCMV source time courses were extracted for each vertex within the ROIs. Within each ROI, we formed feature vectors from vertex-wise source amplitudes at each time point (one vector per trial). Linear support vector machine (SVM) classifiers were trained using data during the encoding phase for each time point. Two kinds of SVM classifiers were trained.

Conceptual classifiers determined whether the picture was a plant or an architecture picture, while perceptual classifiers classified whether the picture was a color or black-and-white picture. Then, we examined the conceptual and perceptual reinstatement using conceptual and perceptual classifiers, which were tested on picture-retrieval trials for each time point. Conceptual and perceptual classification accuracy time series were averaged across vmPFC ROIs and VVC ROIs, and time series were temporally smoothed with a Gaussian kernel (FWHM = 25 ms). We interpret higher cross-phase accuracy as stronger reinstatement of category-specific perceptual and conceptual representations during picture retrieval. SVM classification analysis was performed using the scikit-learn toolbox in Python^68^.

In the group analysis, we exported conceptual and perceptual reinstatement time series for each participant from the cross-phase classification analysis. The analyzed time window was from 0 to 400 ms. Conceptual and perceptual reinstatement strength were contrasted for each time point within the time window using paired t-tests separately for vmPFC and VVC. Cluster-based permutation testing was performed to control for multiple comparisons (cluster-forming p < 0.05; _α_ = 0.05; 2,000 permutations). The results were visualized using the seaborn library^69^. Because we used an automatic cued-recall paradigm, participants heard sounds during both encoding and retrieval, and these trials served as the training and testing sets for the classification analysis. Importantly, the results are unlikely to reflect auditory perception, because sound–picture pairings were randomized across participants. The classification was categorical. Therefore, any given sound item was paired with different perceptual or conceptual picture categories across participants. As a result, group-level differences in classification performance reflect differences in perceptual or conceptual picture category processing rather than sound category processing.

### Frequency-specific encoding-retrieval cross-phase classification analysis

We filtered the broadband source time series from 2 to 40 Hz in 0.5 Hz steps. Then, the same procedure as the encoding-retrieval cross-phase classification analysis was performed on narrowband time series at each frequency. The averaged vmPFC and VVC frequency-specific reinstatement time series were further averaged across four frequency bands: theta (4–8 Hz), alpha (8–13 Hz), beta (13–30 Hz), and gamma (30–40 Hz). The same group analysis comparing conceptual and perceptual reinstatement with cluster-based correction was performed for each frequency band.

### Conceptual and perceptual reinstatement latency comparison

Conceptual and perceptual reinstatement latencies for each participant were defined as the peak latencies within the 0–400 ms range. Paired t-tests were performed to compare the peak latencies between conceptual and perceptual reinstatement.

### Informational cross-correlation analysis

We performed informational cross-correlation analysis to test lead–lag relationships between conceptual and perceptual reinstatement. We computed lagged cross-correlations on individual ROI-averaged conceptual reinstatement time series in the vmPFC and perceptual reinstatement time series in the VVC, using 15 lags (10–150 ms; with 10 ms steps at 100 Hz). A directionality index D(k) was defined at each lag k ^as the difference between r^_conceptual→perceptual_^(k) and r^_perceptual→conceptual_^(k).^ Group inference on D(k) used a two-tailed t-test with permutation testing across subjects with cluster-based correction over the lag dimension (cluster-forming p < 0.05; cluster _α_ = 0.05; 2,000 permutations).

### Spectral state-space Granger causality analysis

We tested directed interactions between vmPFC and VVC during retrieval using spectral state-space Granger causality on time-reversed single-trial source estimates during picture retrieval^70^. Within each ROI, vertex activity was summarized using the mne.extract_label_time_course function with “pca_flip” mode. Spectral time-reversed GC was computed using vmPFC_→_VVC and VVC_→_vmPFC index sets (mne_connectivity.spectral_connectivity_epochs). For each participant, we obtained GC spectra (vmPFC_→_VVC and VVC_→_vmPFC) across 2 to 40 Hz. Similar to the frequency-specific cross-phase classification analysis, we also averaged GC estimates across four frequency bands: theta (4–8 Hz), alpha (8–13 Hz), beta (13–30 Hz), and gamma (30–40 Hz). Then, we performed a paired t-test between GC (vmPFC_→_VVC) and GC (VVC_→_vmPFC) to examine whether the information flow during picture retrieval is from vmPFC to VVC or vice versa.

## Acknowledgments

**Funding**

This work was supported by

## Data and materials availability

The paper and/or the supplementary materials contain all the data needed to evaluate the conclusions. The behavioral and fMRI data that support this study’s findings will be available on OSF.

## Reference

1. Vogt, S. & Magnussen, S. Expertise in Pictorial Perception: Eye-Movement Patterns and Visual Memory in Artists and Laymen. Perception 36, 91–100 (2007).

2. Wing, E. A., Burles, F., Ryan, J. D. & Gilboa, A. The structure of prior knowledge enhances memory in experts by reducing interference. Proc. Natl. Acad. Sci. U.S.A. 119, e2204172119 (2022).

3. Konkle, T., Brady, T. F., Alvarez, G. A. & Oliva, A. Conceptual distinctiveness supports detailed visual long-term memory for real-world objects. Journal of Experimental Psychology: General 139, 558–578 (2010).

4. Josselyn, S. A., Köhler, S. & Frankland, P. W. Finding the engram. Nat Rev Neurosci 16, 521–534 (2015).

5. Josselyn, S. A. & Tonegawa, S. Memory engrams: Recalling the past and imagining the future. Science 367, eaaw4325 (2020).

6. Teyler, T. J. & DiScenna, P. The hippocampal memory indexing theory. Behavioral Neuroscience 100, 147–154 (1986).

7. Hebscher, M., Wing, E., Ryan, J. & Gilboa, A. Rapid Cortical Plasticity Supports Long-Term Memory Formation. Trends in Cognitive Sciences 23, 989–1002 (2019).

8. Moscovitch, M., Cabeza, R., Winocur, G. & Nadel, L. Episodic Memory and Beyond: The Hippocampus and Neocortex in Transformation. Annu. Rev. Psychol. 67, 105–134 (2016).

9. Sekeres, M. J., Winocur, G. & Moscovitch, M. The hippocampus and related neocortical structures in memory transformation. Neuroscience Letters 680, 39–53 (2018).

10. Mesulam, M. M. From sensation to cognition. Brain 121, 1013–1052 (1998).

11. Ranganath, C. & Ritchey, M. Two cortical systems for memory-guided behaviour. Nat Rev Neurosci 13, 713–726 (2012).

12. Bone, M. B., Ahmad, F. & Buchsbaum, B. R. Feature-specific neural reactivation during episodic memory. Nat Commun 11, 1945 (2020).

13. Dijkstra, N., Bosch, S. E. & Gerven, M. A. J. van. Vividness of Visual Imagery Depends on the Neural Overlap with Perception in Visual Areas. J. Neurosci. 37, 1367–1373 (2017).

14. Xue, G. The Neural Representations Underlying Human Episodic Memory. Trends in Cognitive Sciences 22, 544–561 (2018).

15. Wheeler, M. E., Petersen, S. E. & Buckner, R. L. Memory’s echo: Vivid remembering reactivates sensory-specific cortex. Proceedings of the National Academy of Sciences 97, 11125–11129 (2000).

16. Ritchey, M., Wing, E. A., LaBar, K. S. & Cabeza, R. Neural Similarity Between Encoding and Retrieval is Related to Memory Via Hippocampal Interactions. Cerebral Cortex 23, 2818–2828 (2013).

17. Bartlett, F. C. & Kintsch, W. Remembering: A Study in Experimental and Social Psychology. (Cambridge University Press, 1995). doi:10.1017/CBO9780511759185.

18. Schacter, D. L. & Addis, D. R. The cognitive neuroscience of constructive memory: remembering the past and imagining the future. Philosophical Transactions of the Royal Society B: Biological Sciences 362, 773–786 (2007).

19. Schacter, D. L., Norman, K. A. & Koutstaal, W. The cognitive neuroscience of constructive memory. in False-memory creation in children and adults: Theory, research, and implications 129–168 (Lawrence Erlbaum Associates Publishers, Mahwah, NJ, US, 2000).

20. Zhang, L., Alain, C. & Buchsbaum, B. R. Differential reliance on sensory reinstatement and internally transformed representation during vivid retrieval of visual and auditory episodes. Preprint at 10.1101/2025.06.16.659408 (2025).

21. Gordon, A. M., Rissman, J., Kiani, R. & Wagner, A. D. Cortical Reinstatement Mediates the Relationship Between Content-Specific Encoding Activity and Subsequent Recollection Decisions. Cereb Cortex 24, 3350–3364 (2014).

22. Barker, R. M., St-Laurent, M. & Buchsbaum, B. R. Neural reactivation and judgements of vividness reveal separable contributions to mnemonic representation. NeuroImage 255, 119205 (2022).

23. Heinbockel, H., Wagner, A. D. & Schwabe, L. Post-retrieval stress impairs subsequent memory depending on hippocampal memory trace reinstatement during reactivation. Science Advances 10, eadm7504 (2024).

24. Bowman, C. R. & Zeithamova, D. Abstract Memory Representations in the Ventromedial Prefrontal Cortex and Hippocampus Support Concept Generalization. J. Neurosci. 38, 2605–2614 (2018).

25. Ghosh, V. E., Moscovitch, M., Colella, B. M. & Gilboa, A. Schema Representation in Patients with Ventromedial PFC Lesions. J. Neurosci. 34, 12057–12070 (2014).

26. Giuliano, A. E., Bonasia, K., Ghosh, V. E., Moscovitch, M. & Gilboa, A. Differential Influence of Ventromedial Prefrontal Cortex Lesions on Neural Representations of Schema and Semantic Category Knowledge. J Cogn Neurosci 33, 1928–1955 (2021).

27. Kerrén, C., Reznik, D., Doeller, C. F. & Griffiths, B. J. Exploring the role of dimensionality transformation in episodic memory. Trends in Cognitive Sciences 29, 614–626 (2025).

28. Mack, M. L., Preston, A. R. & Love, B. C. Ventromedial prefrontal cortex compression during concept learning. Nat Commun 11, 46 (2020).

29. Bein, O. & Niv, Y. Schemas, reinforcement learning and the medial prefrontal cortex. Nat. Rev. Neurosci. 26, 141–157 (2025).

30. Lifanov, J., Linde-Domingo, J. & Wimber, M. Feature-specific reaction times reveal a semanticisation of memories over time and with repeated remembering. Nat Commun 12, 3177 (2021).

31. Linde-Domingo, J., Treder, M. S., Kerrén, C. & Wimber, M. Evidence that neural information flow is reversed between object perception and object reconstruction from memory. Nat Commun 10, 179 (2019).

32. Guth, T. A. et al. Theta-phase locking of single neurons during human spatial memory. Nat Commun 16, 7402 (2025).

33. Herweg, N. A., Solomon, E. A. & Kahana, M. J. Theta Oscillations in Human Memory. Trends in Cognitive Sciences 24, 208–227 (2020).

34. Kota, S., Rugg, M. D. & Lega, B. C. Hippocampal Theta Oscillations Support Successful Associative Memory Formation. J. Neurosci. 40, 9507–9518 (2020).

35. Rudoler, J. H., Herweg, N. A. & Kahana, M. J. Hippocampal Theta and Episodic Memory. J. Neurosci. 43, 613–620 (2023).

36. Ursino, M. & Pirazzini, G. Theta–gamma coupling as a ubiquitous brain mechanism: implications for memory, attention, dreaming, imagination, and consciousness. Current Opinion in Behavioral Sciences 59, 101433 (2024).

37. Anderson, K. L., Rajagovindan, R., Ghacibeh, G. A., Meador, K. J. & Ding, M. Theta Oscillations Mediate Interaction between Prefrontal Cortex and Medial Temporal Lobe in Human Memory. Cereb Cortex 20, 1604–1612 (2010).

38. Backus, A. R., Schoffelen, J.-M., Szebényi, S., Hanslmayr, S. & Doeller, C. F. Hippocampal-Prefrontal Theta Oscillations Support Memory Integration. Current Biology 26, 450–457 (2016).

39. Hebscher, M., Meltzer, J. A. & Gilboa, A. A causal role for the precuneus in network-wide theta and gamma oscillatory activity during complex memory retrieval. eLife 8, e43114 (2019).

40. Barry, D. N., Barnes, G. R., Clark, I. A. & Maguire, E. A. The Neural Dynamics of Novel Scene Imagery. J. Neurosci. 39, 4375–4386 (2019).

41. Fuentemilla, L., Barnes, G. R., Düzel, E. & Levine, B. Theta oscillations orchestrate medial temporal lobe and neocortex in remembering autobiographical memories. NeuroImage 85, 730–737 (2014).

42. Kaplan, R. et al. Medial prefrontal theta phase coupling during spatial memory retrieval. Hippocampus 24, 656–665 (2014).

43. Barron, H. C. et al. Unmasking Latent Inhibitory Connections in Human Cortex to Reveal Dormant Cortical Memories. Neuron 90, 191–203 (2016).

44. Tomita, H., Ohbayashi, M., Nakahara, K., Hasegawa, I. & Miyashita, Y. Top-down signal from prefrontal cortex in executive control of memory retrieval. Nature 401, 699–703 (1999).

45. Guise, K. G. & Shapiro, M. L. Medial Prefrontal Cortex Reduces Memory Interference by Modifying Hippocampal Encoding. Neuron 94, 183–192.e8 (2017).

46. McCormick, C., Barry, D. N., Jafarian, A., Barnes, G. R. & Maguire, E. A. vmPFC Drives Hippocampal Processing during Autobiographical Memory Recall Regardless of Remoteness. Cereb Cortex 30, 5972–5987 (2020).

47. Robin, J. & Moscovitch, M. Details, gist and schema: Hippocampal–neocortical interactions underlying recent and remote episodic and spatial memory. Current Opinion in Behavioral Sciences 17, 114–123 (2017).

48. Mesulam, M.-M. Behavioral Neuroanatomy Large-Scale Networks, Association Cortex, Frontal Syndromes, the Limbic System, and Hemispheric Specializations. in *Principles of Behavioral and Cognitive Neurology* (ed. Mesulam, M.-M.) 0 (Oxford University Press, 2000). doi:10.1093/oso/9780195134759.003.0001.

49. Winocur, G. & Moscovitch, M. Memory Transformation and Systems Consolidation. Journal of the International Neuropsychological Society 17, 766–780 (2011).

50. Favila, S. E., Lee, H. & Kuhl, B. A. Transforming the Concept of Memory Reactivation. Trends in Neurosciences 43, 939–950 (2020).

51. Gloede, M. E. & Gregg, M. K. The fidelity of visual and auditory memory. Psychon Bull Rev 26, 1325–1332 (2019).

52. Clarke, A., Taylor, K. I., Devereux, B., Randall, B. & Tyler, L. K. From Perception to Conception: How Meaningful Objects Are Processed over Time. Cereb Cortex 23, 187–197 (2013).

53. Gilboa, A. & Moscovitch, M. Ventromedial prefrontal cortex generates pre-stimulus theta coherence desynchronization: A schema instantiation hypothesis. Cortex 87, 16–30 (2017).

54. Griffiths, B. J. & Jensen, O. Gamma oscillations and episodic memory. Trends in Neurosciences 46, 832–846 (2023).

55. Mysin, I. & Shubina, L. From mechanisms to functions: The role of theta and gamma coherence in the intrahippocampal circuits. Hippocampus 32, 342–358 (2022).

56. Min Park, Y., Park, J., Young Kim, I., Koo Kang, J. & Pyo Jang, D. Interhemispheric theta coherence in the hippocampus for successful object-location memory in human intracranial encephalography. Neuroscience Letters 786, 136769 (2022).

57. Nyhus, E. & Curran, T. Functional role of gamma and theta oscillations in episodic memory. Neuroscience & Biobehavioral Reviews 34, 1023–1035 (2010).

58. Welle, C. G. & Contreras, D. Sensory-driven and spontaneous gamma oscillations engage distinct cortical circuitry. J Neurophysiol 115, 1821–1835 (2016).

59. Wilming, N., Murphy, P. R., Meyniel, F. & Donner, T. H. Large-scale dynamics of perceptual decision information across human cortex. Nat Commun 11, 5109 (2020).

60. Lizarazu, M., Lallier, M. & Molinaro, N. Phase−amplitude coupling between theta and gamma oscillations adapts to speech rate. Annals of the New York Academy of Sciences 1453, 140–152 (2019).

61. Busch, N. A., Dubois, J. & VanRullen, R. The Phase of Ongoing EEG Oscillations Predicts Visual Perception. J. Neurosci. 29, 7869–7876 (2009).

62. Michalareas, G. et al. Alpha-Beta and Gamma Rhythms Subserve Feedback and Feedforward Influences among Human Visual Cortical Areas. Neuron 89, 384–397 (2016).

63. Bastos, A. M. et al. Canonical Microcircuits for Predictive Coding. Neuron 76, 695–711 (2012).

64. Faul, F., Erdfelder, E., Lang, A.-G. & Buchner, A. G*Power 3: a flexible statistical power analysis program for the social, behavioral, and biomedical sciences. Behav Res Methods 39, 175–191 (2007).

65. Larson, E., et al. MNE-Python. Zenodo 10.5281/ZENODO.592483 (2025).

66. Fischl, B. FreeSurfer. NeuroImage 62, 774–781 (2012).

67. Destrieux, C., Fischl, B., Dale, A. & Halgren, E. Automatic parcellation of human cortical gyri and sulci using standard anatomical nomenclature. NeuroImage 53, 1–15 (2010).

68. Pedregosa, F. et al. Scikit-learn: Machine Learning in Python. J. Mach. Learn. Res. 12, 2825–2830 (2011).

69. Waskom, M.. seaborn: statistical data visualization. JOSS 6, 3021 (2021).

70. Winkler, I., Panknin, D., Bartz, D., Muller, K.-R. & Haufe, S. Validity of Time Reversal for Testing Granger Causality. IEEE Trans. Signal Process. 64, 2746–2760 (2016).

