## Supplementary material for "Theta-mediated conceptual reinstatement in vmPFC precedes perceptual reinstatement in ventral visual cortex during memory recall": Figure S1

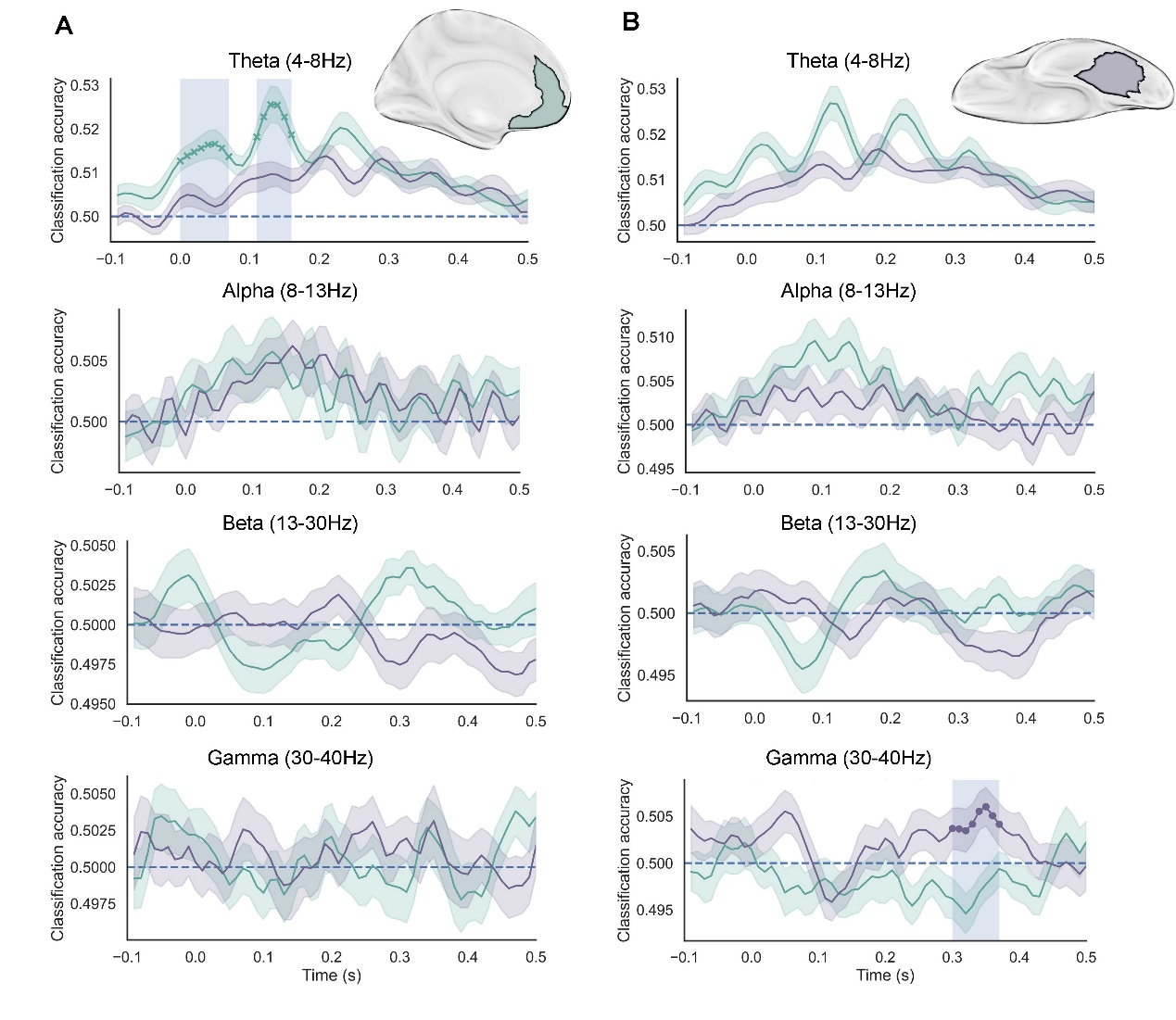

Figure S1. **Conceptual advantage in vmPFC expressed in theta band while leads perceptual advantage in VVC expressed in gamma band.** (A). In vmPFC, conceptual reinstatement was significantly stronger than perceptual reinstatement in the theta band during the two time windows: 0–70 ms and 110–160 ms ($p_{fwe}$ < 0.05). No significant differences were observed in other frequency bands. (B) In contrast, VVC showed significantly stronger perceptual than conceptual reinstatement in the gamma band from 300–370 ms ($p_{fwe}$ < 0.05). No significant differences were observed in other frequency bands. vmPFC, ventromedial prefrontal cortex; VVC, ventral visual cortex.
